# Platform for Identifying Human Glycan-Specific Antibodies Against Bacterial Pathogens using Synthetic Glycan Fragments

**DOI:** 10.1101/2025.08.19.671086

**Authors:** A. Robin Temming, Mathieu Claireaux, Gius Kerster, Silvie E. Groenewege, Thijs Voskuilen, Zhen Wang, Jeroen D.C. Codée, Marit J. van Gils, Nina M. van Sorge

**Author notes:** authors contributed equally (Mathieu Claireaux, Gius Kerster).

## Abstract

Bacterial infections represent a substantial global health challenge, impacting both human and veterinary health. The ongoing evolution of antibiotic-resistant pathogens, coupled with limited new antibiotic discoveries, urges the need for alternative strategies to treat and prevent these infections. Passive immunization with monoclonal antibodies (mAbs) is gaining interest as a promising alternative. Here, we report an experimental pipeline for generating human mAbs from healthy donor B cells using synthetic mimics of complex bacterial glycans. We identified functional mAbs recognizing discrete and unique epitopes on the surface glycans of two bacterial priority pathogens; *Staphylococcus aureus* and *Streptococcus pyogenes*. The use of chemically-defined synthetic glycans was critical for the discovery and systematic characterization of mAbs. From a heterogeneous mix of B cell specificities, mAbs were isolated with reactivities against immunodominant but also to less common or even masked epitopes. The pipeline can be adapted to different glycan targets, donor material or specific antibody isotypes. This work thereby paves the way for the discovery of glycan-specific mAbs with clinical relevance to treat, prevent or diagnose infections with *S. aureus*, *S. pyogenes* or other bacterial pathogens.

## INTRODUCTION

Bacterial infections represent a global burden for human and veterinary health with persistent mortality and morbidity. Antibiotics have long been considered, and continue to be, the gold standard for treating bacterial infections. However, the effectiveness of antibiotics has declined over time due to the emergence of drug-resistant strains, known as antimicrobial resistance (AMR), which pose a significant global health threat. The World Health Organization (WHO) has recognized AMR as a global health crisis, resulting in studies that identified priority pathogens and priority areas of vaccine development (1–3). The ongoing evolution of antibiotic-resistant pathogens, coupled with limited antibiotic discoveries, urgently calls for alternative strategies to treat and prevent of bacterial infections.

Boosting host immunity to infectious pathogens, e.g. passive immunization with antibacterial antibodies, represents a promising alternative to antibiotics for prevention and possibly treatment (4). Antibodies can confer protective effects through several mechanisms, including neutralizing bacterial toxins or facilitating bacterial opsonization to enhance phagocytic uptake and killing. Administration of monoclonal antibodies (mAbs) provides several features advantageous or complementary to active immunization, i.e. it offers immediate effects, is effective in immunocompromised individuals, and provides transient immunity instead of long-lasting, thereby exerting less evolutionary pressure on pathogens to develop evasion strategies. Therapeutic application of mAbs may be beneficial in established infections, or prophylactically for individuals at risk, e.g. scheduled hospital procedures. For both therapeutic or prophylactic use of mAbs, the choice of the target antigen is critical in terms of accessibility, abundance, conservation, and uniqueness, i.e. limited cross-reactivity to other bacteria or host molecules.

*Staphylococcus aureus* (*S. aureus*) is one of the high-priority pathogens on the WHO list (1). This Gram-positive opportunistic pathogen is a leading cause of hospital- and community-acquired infections, including skin infections and bacteremia (5, 6). Moreover, deaths associated or attributable to the antibiotic-resistant variant methicillin-resistant *S. aureus* (MRSA) more than doubled between 1990 and 2021, rendering MRSA the species with the greatest health loss worldwide in 2021 (2). These numbers underscore the need for further research for alternative prevention and treatment strategies, such as antibody-based strategies.

The surface of *S. aureus* is extensively decorated with glycopolymers called wall teichoic acid (WTA) that are accessible and abundant targets for human antibodies (7–10). WTA expression is conserved across *S. aureus* strains and is an immunodominant antigen as reflected by the ubiquitous presence of anti-WTA antibodies in the sera of virtually all tested healthy individuals (9–11). In addition, WTA has been identified as a critical target antigen in the host defense against *S. aureus* (8, 12). In the large majority of *S. aureus* strains, WTA consists of polymerized ribitol phosphate (RboP) units with D-alanine and *N-*acetylglucosamine (GlcNAc) side groups, which exhibit limited antigenic variation. The variation in GlcNAc depends on the presence and expression levels of dedicated Tar glycosyltransferases (11, 13, 14). *tarS* is part of *S. aureus*’ core genome and encodes for TarS, which attaches GlcNAc on C4 of RboP in β configuration (β-1,4-GlcNAc). Approximately 37% and 7% of the *S. aureus* population co-encode additional *tar* glycosyltransferases *tarM* and *tarP*, which attach GlcNAc to C4 in α configuration (α-1,4-GlcNAc) and to C3 in β configuration, respectively (11). The resulting limited glycovariation in WTA has implications in immune responses and mAb-based strategies targeting WTA.

Natural exposure to *S. aureus* elicits WTA-specific IgG antibodies that predominantly target β-GlcNAc epitopes, with no cross-reactivity to α-GlcNAc. WTA-specific mAbs targeting β-GlcNAc or α-GlcNAc epitopes exist and have been previously isolated from memory and plasma B cells derived from *S. aureus* patients during the convalescent phase (15). These mAbs, expressed as IgG1 and conjugated to antibiotics, have been proven effective against *S. aureus* in mouse infection models by reducing bacterial persistence (8, 15). Nevertheless, so far, the workflow leading to the WTA-specific mAbs was extremely inefficient and labor-intensive. Because naturally-sourced WTA (and isolated bacterial glycans in general) being heterogeneous in size and modifications, lacking a chemical handle to orient the molecule, and being instable after isolation resulting in loss of epitopes, previous approaches were not able to preselect WTA-specific B cells and therefore relied on large screening of cultured single B cell supernatants. To improve future discovery efforts, we developed a pipeline based on single-cell sorting of glycan-specific B cells using synthetic WTA fragments as stable antigen probes in combination with RNA sequencing of antibody variable domains. Building on an existing strategy developed for protein-based antigens (successfully applied to identify anti-viral mabs e.g. anti-SARS-CoV-2 spike mAbs (16)), we report an optimized scBCRseq workflow for identifying functional mAbs targeting different glycan epitopes on the *S. aureus* WTA. We show the versatility of this workflow by generating mAbs against surface glycopolymers of another human priority pathogens, i.e. *Streptococcus pyogenes* (Group A streptococcus; GAS). Taken together, this study demonstrates the utility of synthetic mimics of bacterial glycans in facilitating the discovery of mAbs against complex natural glycan antigens.

## RESULTS

### Design and verification of synthetic WTA probes

We designed fluorescently labeled synthetic WTA (sWTA) probes to identify and sort human memory B cells specific to WTA,. These probes consisted of biotinylated RboP hexamers enzymatically modified with GlcNAc side groups through the activity of the specific Tar enzymes. (Figure 1A). These sWTA structures are stable, short fragments modified with either β-1,4-, β-1,3-, or α-1,4-GlcNAc decorations, instead of the more complex, heterogeneous, and unstable natural WTA that can be isolated from *S. aureus*. The sWTA structures have been previously characterized and allowed for the detection of antibodies in pooled human serum and patients with *S. aureus* bacteremia (9, 10). The sWTA probes were coupled to fluorophore-conjugated streptavidin andtested for their specificity and ability to engage previously-identified WTA-specific mAbs (against β- or α-GlcNAc) coated on beads as surrogates for WTA-specific B cells using flow cytometry. The efficient glycoform-specific detection of anti-WTA coated beads and not isotype control (Figure 1B) indicated that these sWTA probes were suitable to detect WTA-reactive B cells in the total B cell pool.

**Figure 1:**
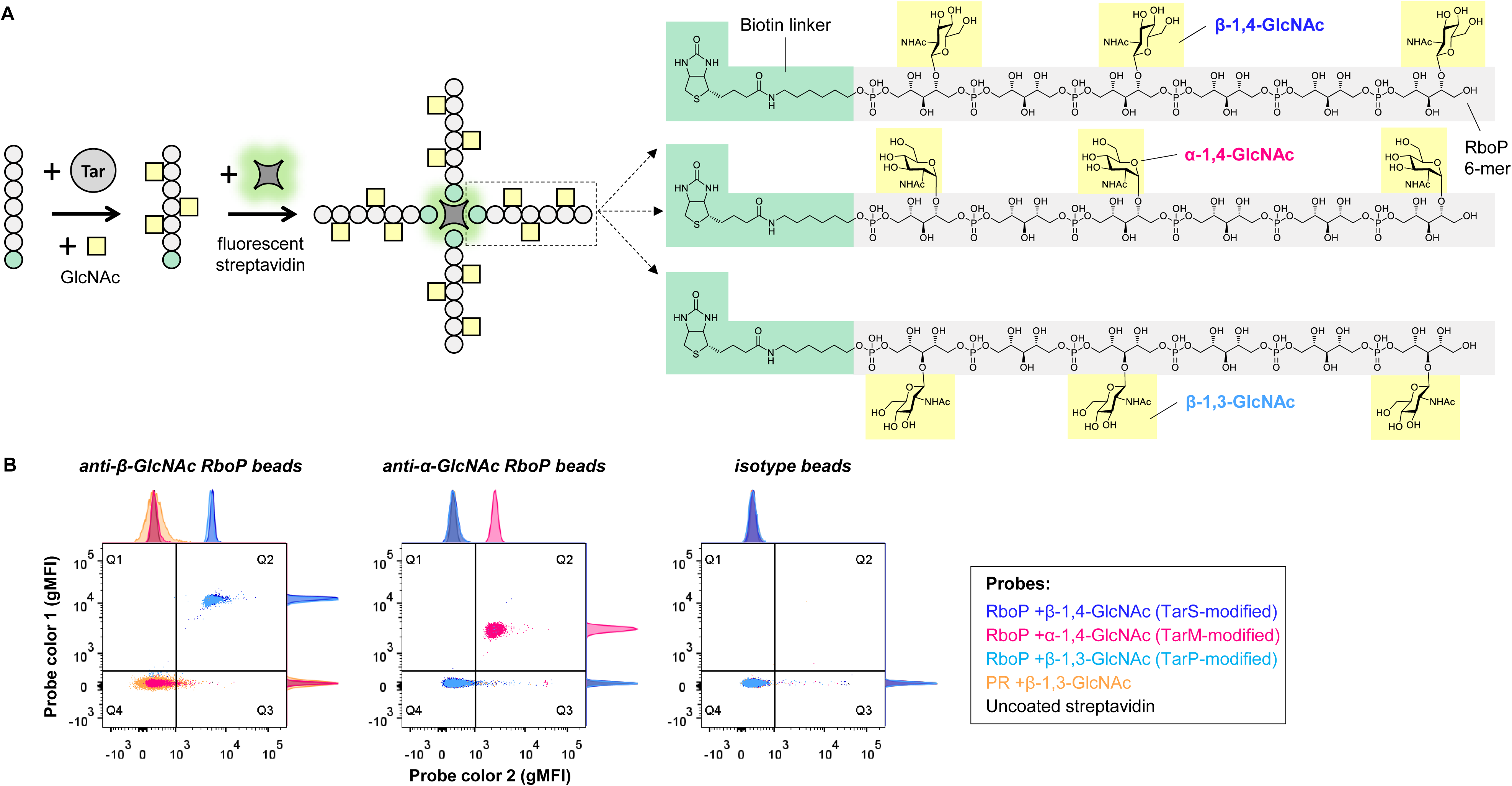
sWTA probe generation and validation. **A** Schematic representation of sWTA probe generation for all three glycoforms, i.e. RboP +β-1,4-GlcNAc, +α-1,4-GlcNAc, or +β-1,3-GlcNAc. All sWTA probes were made for detection in two different fluorescence channels using streptavidin conjugated to AF647 or BB515. **B** Dual sWTA probe labeling of protein A-coated beads coated with anti-β-GlcNAc RboP (clone 4497), anti-α-GlcNAc RboP (clone 4461), and isotype IgG1. Data in dot plots represent geometric mean fluorescence intensities (gMFI) signals (fluorophores: AF647 and BB515) on the beads. Q2 and Q4 comprise, respectively, double positive (dual sWTA probe labeling) and double negative (no sWTA probe binding) beads. Signals within Q1 and Q3 represent aspecific binding of, respectively, AF647 and BB515 streptavidin to beads. Histograms are included on the sides of the dot plots to visualize relative amounts of different bead populations within a fluorescent channel.

A large buffy coat sample from a healthy donor (HD), yielding 376 × 10^6^ PBMCs, was used to isolate *S. aureus* WTA-specific memory B cells. Before sorting, we screened buffy coat plasma for the presence of antibodies against the different WTA-GlcNAc glycoforms using our previously published WTA bead assay (9, 10) (Figure S1). Antibody profiles demonstrated that all WTA reactivities of interest are distinctly present for plasma IgG, IgM and IgA. The abundant presence of anti-WTA antibodies suggests that this material is suitable as a source to identify and sort WTA-specific memory B cells.

### sWTA probes identify WTA-specific memory B cells

A B cell staining approach using dual-labeled antigenic probes was implemented to improve detection of WTA-specific B cells while minimizing false positive selection of fluorophore-specific B cells. To this end, each of the three sWTA glycoform-specific probes (i.e. RboP hexamers + β-1,4-, α-1,4-, or β-1,3-GlcNAc) was conjugated to the same two fluorophores to ensure reliable detection of all specificities. B cells were first enriched from PBMCs using negative selection and subsequently stained with a mixture of the three dual-labeled sWTA glycoform probes. WTA-specific B cells were detected by flow cytometry and represented 0.004% of the live CD19+ B cell fraction in this particular healthy donor (Figure S2A). A total of 78 WTA-specific B cells were single cell sorted from this modest fraction and further characterized based on their IgD and CD27 surface expression defining naive and memory B cell populations. Notably, the majority (73 of 78; 94%) of the probe-labeled sorted B cells exhibited a CD27+ memory B cell phenotype (Figure S2B), confirming prior exposure to WTA antigens and potential role in immune memory response. Most (54 of 73; 74%) of these sorted memory B cells were also positive for surface IgD (Figure S2B), corresponding to an unswitched memory phenotype. The remaining (19 of 73; 26%) sorted B cells were CD27+ IgD−, representing switched memory B cells (Figure S2B).

### B cell-derived mAbs recognize bead-immobilized sWTA

For all sorted sWTA-reactive B cells, the variable heavy (VH) and light (VL) chain sequences were amplified by nested PCR and cloned into expression vectors. Among 78 clones, 21 had matching VH-VL pairs (26.9%), of which 16 were unique (76.2%). Next, 15 of the 16 unique clones were successfully produced as a human IgG1 at pilot scale (Figure S3) and tested for binding to sWTA immobilized onto beads. Reactivity to sWTA was confirmed for 10 out of 15 unique clones (66.7%) (Figure 2). At glycoform level, seven (out of 10; 70%) of these clones bound specifically to β-GlcNAc decorated RboP hexamers, whereas three clones recognized the α-1,4-GlcNAc stereochemical WTA structures. There was no cross-reactivity of mAbs for the other stereochemical epitope of GlcNAc. Notably, clones W1F10, W1C4, and W1B9 showed some distinction between β-1,3- and β-1,4-GlcNAc epitopes, with a preference for β-1,3-GlcNAc over β-1,4-GlcNAc (Figure 2). Index sorting (linking sWTA reactivity to the corresponding sorted B cell) of all VH-VL pair matching clones showed that the sWTA-reactive clones mainly originate from B cells belonging to the memory compartment (low IgD:CD27 ratios), while B cells encoding the unreactive or very low affinity clones generally displayed more naïve phenotypes (high IgD:CD27 ratios) (Figure S4). This suggests that substantial WTA reactivity depends on prior engagement of B cells with the antigen, rather than arising from cross-reactive naïve B cells. Next, sWTA-reactive clones were produced at larger scale to compare binding capacities at equimolar level (Figure 3). The three α-GlcNAc binders showed the highest and comparable binding efficiency (W1C11 = W1E11 > W1B6; Figure 3A). In contrast, there was substantial variation in binding capacity among β-GlcNAc-specific mAbs with W1F10 and W1G7 being the most efficient clones (Figure 3B). The lack of binding to β-1,3-GlcNAc attached to a chemically-synthesized polyrhamnose (PR) hexamer (as expressed by *S. pyogenes*) demonstrated that the mAb clones were specific for *S. aureus* WTA-GlcNAc and that recognition depended on the attachment of GlcNAc to the RboP backbone (Figure 3B).

**Figure 2:**
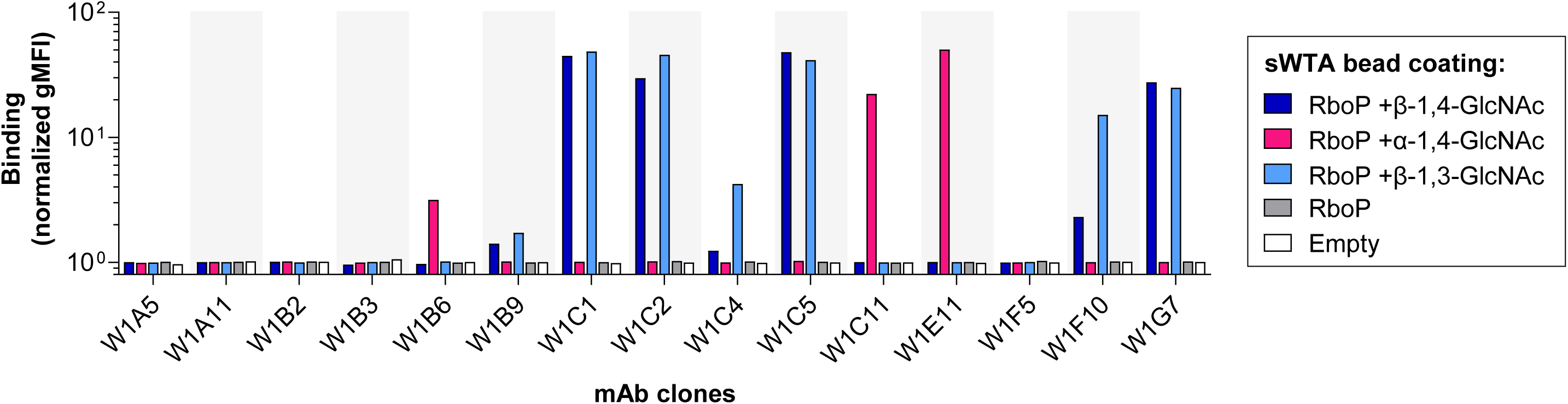
sWTA specificity screening of pilot-scale produced mAbs. Binding profiles of 15 B cell-derived mAbs expressed by HEK293T cells (production levels in **Figure S3**) as human IgG1 to streptavidin-coated beads immobilized with biotinylated sWTA to determine clone reactivity as measured by flow cytometry. Fluorescent signals are depicted in this figure as gMFI fold changes relative to the condition without antibodies to compensate for technical variation. Empty beads were used to determine background.

**Figure 3:**
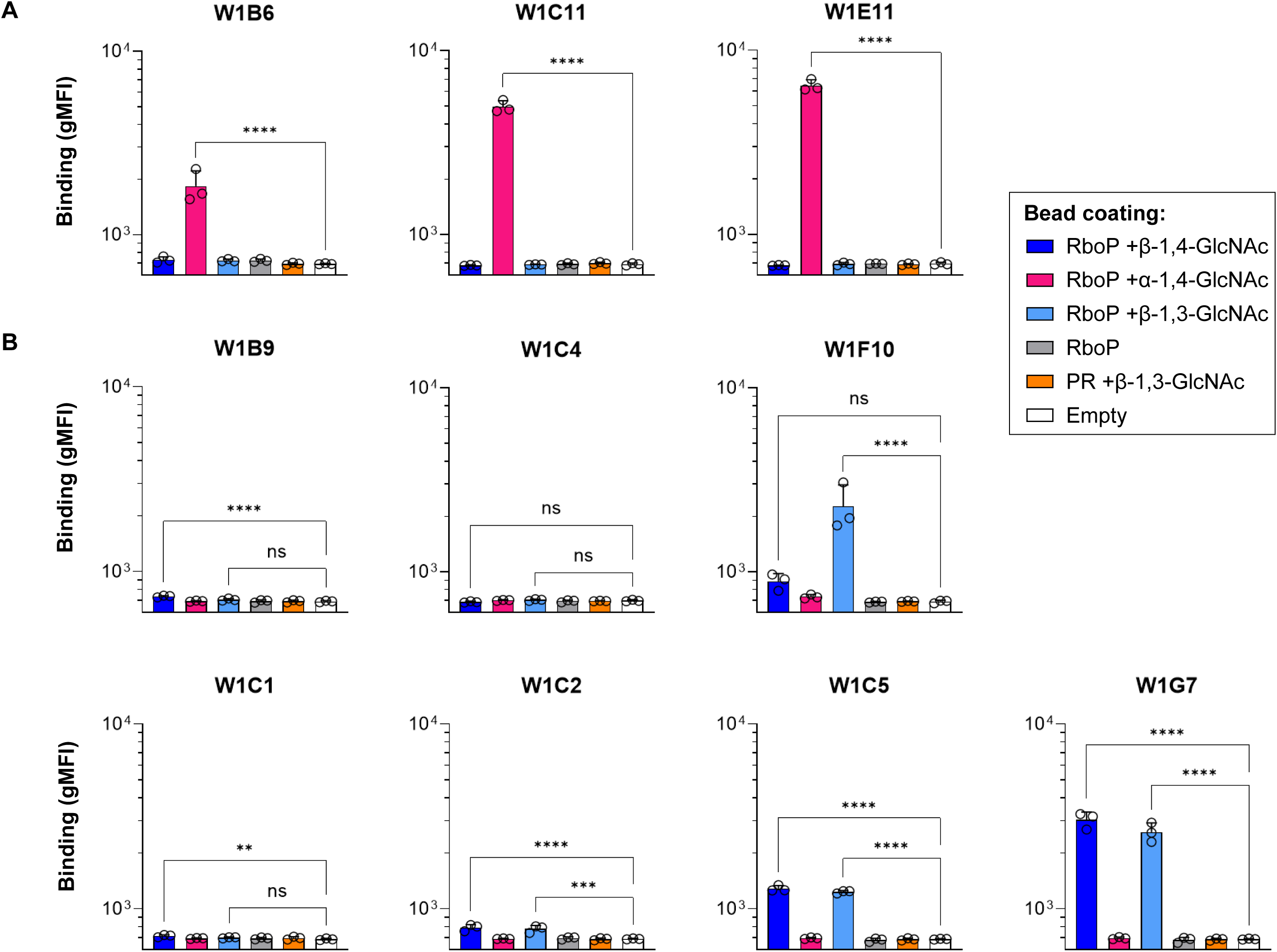
Specificity verification of sWTA-reactive mAbs at equimolar level. Clones that displayed sWTA reactivity in the pilot screening (**main** Figure 2) were selected for large-scale production in HEK293 Freestyle cells and purified through protein G agarose. Selected clones were categorized based on their α-GlcNAc (**A**) or β-GlcNAc (**B**) reactivity and relative binding capacities to sWTA beads were determined at a concentration of 3 μg/ml. Beads coated with polyrhamnose (PR) +β-1,3-GlcNAc and empty beads were used as controls for cross-reactivity and background, respectively. IgG1 binding to sWTA beads was measured by flow cytometry and data represent the gMFI mean ± s.d. of three independent experiments. One-way ANOVA with Dunnett’s multiple comparisons test was performed to determine significant binding of sWTA-reactive clones to glycan-coated beads compared to empty beads. ns not significant ** p < 0.01 **** p < 0.0001. Index sort data of all sorted clones can be found in Figure S4.

### B cell-derived mAbs recognize naturally-expressed WTA on *S. aureus*

We next determined whether the identified clones were able to recognize the more complex WTA structures expressed on bacterial surfaces that differ in glycopolymer length, heterogeneity, presence of D-alanine side groups and attachment to the thick peptidoglycan layer. For each glycoform specificity, one mAb clone was selected based on evident binding in the bead assay. *S. aureus* strains (lacking IgG-binding protein A [Δ*spa*]) were incubated with a concentration range of mAb to assess binding. β-GlcNAc specific clones W1G7 (β-1,3 = β-1,4-GlcNAc) and W1F10 (β-1,3 > β-1,4-GlcNAc) showed binding (areas under the curve [AUC] of 3,549 and 499, respectively) to strain N315 that expresses β-1,4-/β-1,3-GlcNAc WTA (Figure 4A). Deletion of both *tar* genes (Δ*tarSP*) completely abolished binding (AUC 6 and 9, respectively; Figure 4B). Binding to α-GlcNAc WTA was studied using *S. aureus* strain Newman, which expresses TarM and TarS, resulting in expression of WTA decorated with α-1,4- and β-1,4-GlcNAc, respectively. Clone W1C11 showed concentration-dependent binding to the Newman strain (AUC 15,249; Figure 4C), but not to β-GlcNAc-expressing strain N315 (AUC 11; Figure 4A). Both β-GlcNAc specific clones W1G7 and W1F10 also showed binding to the Newman strain (AUC 10,037 and 123, respectively; Figure 4C), although W1F10 bound to a lower extend compared to N315 likely due to the absence of β-1,3-GlcNAc. Consistent with results from the bead assay, none of the mAbs bound to *S. pyogenes* 5448 Δ*gacH* (AUC 11-16), which expresses PR decorated with β-1,3-GlcNAc but lacks the GlcNAc-capping glycerol phosphate modifications (17) (Figure 4D). Together, these data confirm the ability of the clones to specifically recognize natural WTA via the respective GlcNAc modifications. This approach thereby demonstrates that chemically synthesized glycan mimics of WTA can be used as probes to discover and generate mAbs with reactivities against more complex natural equivalents expressed on bacterial surfaces.

**Figure 4:**
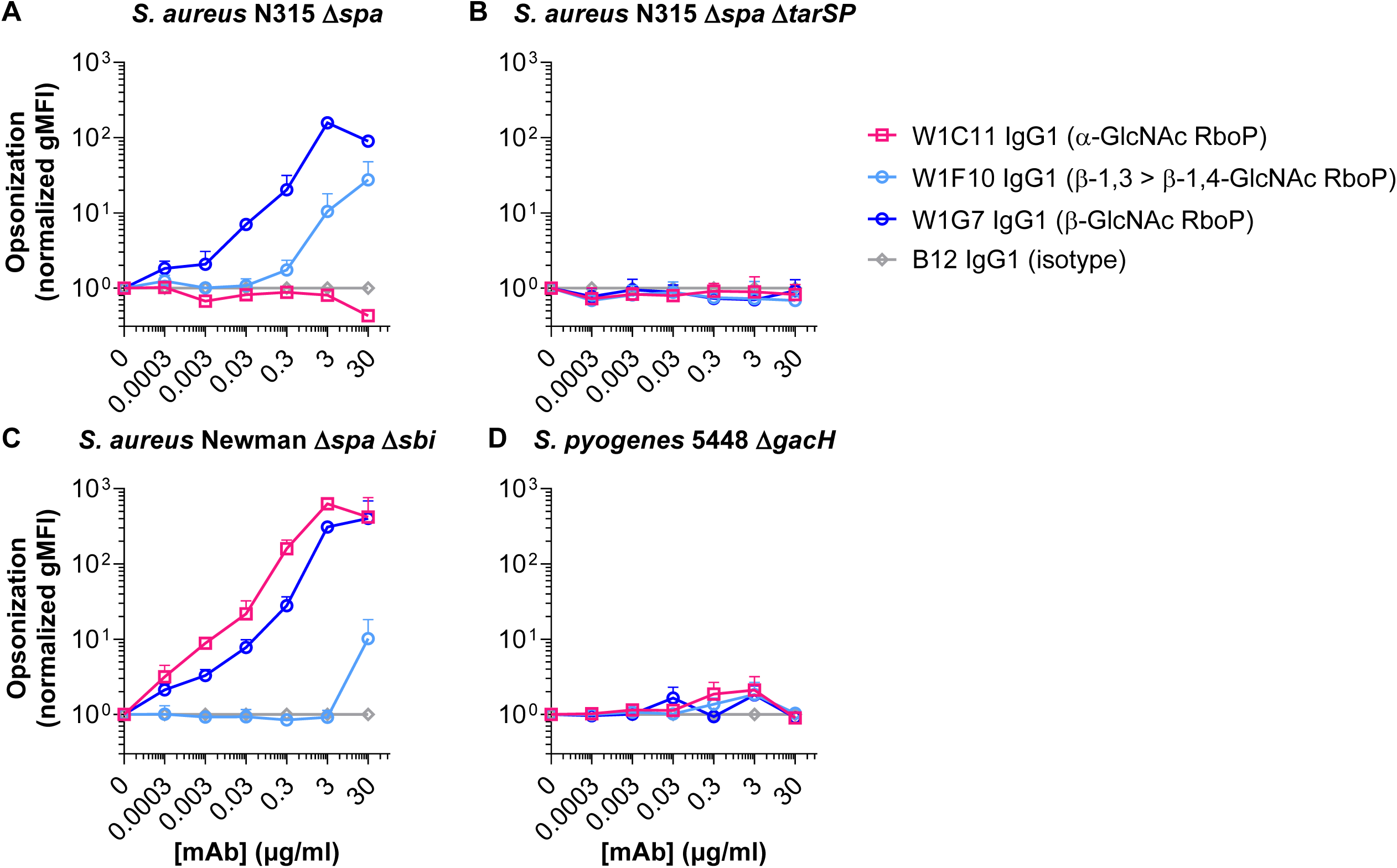
Binding of sWTA-reactive mAbs to WTA on *S. aureus* surface. For each glycoform specificity, one mAb clone was selected to assess bacterial opsonization. **A-D** Binding of W1C11 (anti-α-GlcNAc), W1F10 (anti-β-GlcNAc, with preference for β-1,3-GlcNAc), W1G7 (anti-β-GlcNAc), and B12 (isotype control) to *S. aureus* strains N315 Δ*spa* (**A**), N315 Δ*spa* Δ*tarSP* (**B**), Newman Δ*spa* Δ*sbi* (**C**), and *S. pyogenes* strain 5448 Δ*gacH* (**D**). IgG1 binding to bacteria was measured using flow cytometry and data represent normalized mean gMFI + s.d. (isotype signals set to 1) of three independent experiments. N315 Δ*spa* Δ*tarSP* and *S. pyogenes* 5448 Δ*gacH* were included as controls for WTA GlcNAc (species) specificity.

### The anti-WTA mAbs elicit complement deposition and phagocytosis

To study the functionality of the newly discovered mAbs, we assessed deposition of complement component C3b on *S. aureus’* surface. *S. aureus* strains N315 and Newman were incubated consecutively with W1C11, W1F10, and W1G7 IgG1 and IgG-/IgM-depleted human serum as complement source. The β-GlcNAc binders W1G7 and W1F10 (β-1,3 > β-1,4) induced efficient C3b deposition on strain N315 (AUC 751 and 959, respectively; Figure 5A), which was abolished in the absence of GlcNAc modifications Figure 5B; (Δ*tarSP*; AUC 9 and 3, respectively). In line with binding levels (Figure 4A,C), α-GlcNAc mAb W1C11 IgG1 was inert towards N315 (AUC 5), but readily triggered concentration-dependent C3b deposition on the Newman strain (AUC 1,263; Figure 5C). Also consistent with binding, W1G7 and W1F10 elicited efficient (AUC 828) and negligible (AUC 3) C3b deposition onto Newman, respectively (Figure 5C).

**Figure 5:**
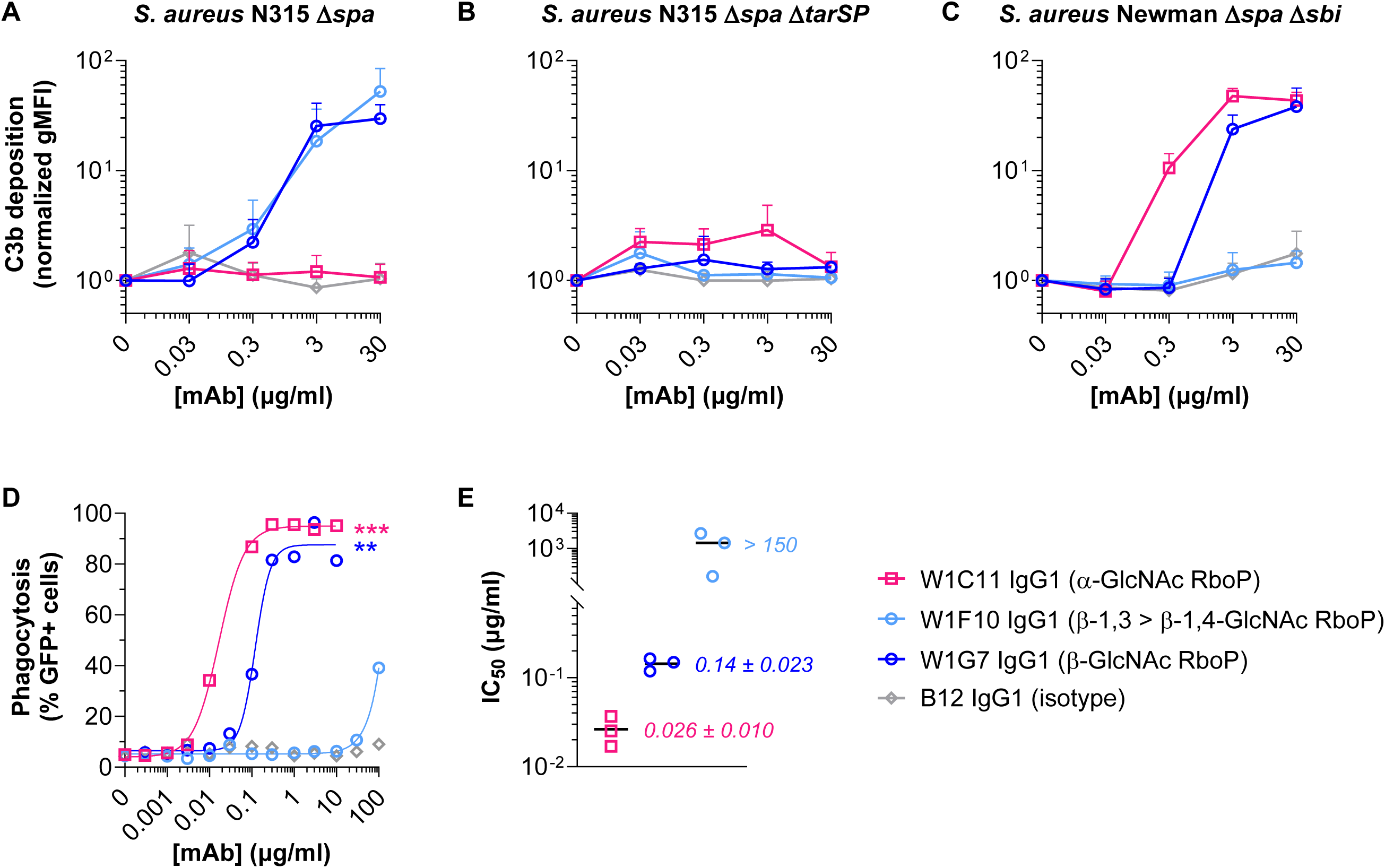
Effector functions of anti-WTA clones towards *S. aureus*. **A-C** C3b deposition by sWTA-reactive mAbs onto *S. aureus* strains N315 Δ*spa* (**A**), N315 Δ*spa* Δ*tarSP* (**B**), and Newman Δ*spa* Δ*sbi* (**C**). Data represent C3b binding (normalized gMFI + s.d.) of three independent experiments as measured by flow cytometry. Fluorescent signals are depicted as a fold change relative to the condition without antibodies to compensate for variation in background signals between biological replicates. **D** Neutrophil-mediated phagocytosis of GFP-expressing *S. aureus* Newman *Δspa Δsbi* by anti-WTA mAbs. Displayed data represent percentages of GFP-positive neutrophils and are representative of three biological replicates (individual replicates can be found in **Figure S5**). Curves were generated using nonlinear dose-response fitting model. **E** Relative phagocytic capacities of the anti-WTA mAbs. Absolute IC_50_ values were determined for each replicate individually using nonlinear dose-response fitting model. Black lines represent means of the IC_50_ values which are depicted as data points. Assays (**A-D**) were performed in the presence of 1% IgG-IgM-depleted human serum as complement source. Statistical differences compared to isotype were determined by One-way ANOVA. ** p < 0.01 *** p < 0.001.

Phagocytosis was investigated as another qualitative assessment of the WTA-specific mAbs. Human neutrophils were incubated with pre-opsonized GFP-expressing *S. aureus* Newman (Δ*spa*/*sbi*) in the presence of IgG-/IgM-depleted human serum as complement source. Background phagocytosis levels with an isotype control antibody fluctuated between 4-10% and represent the fraction of phagocytosis that occurs independent of antibody opsonization. The α-GlcNAc binder W1C11 and β-GlcNAc binder W1G7 IgG1 induced *S. aureus* uptake by nearly all neutrophils (Figure 5D and S5) with an IC_50_ of respectively 0.026 ± 0.01 µg/ml (0.18 ± 0.07 nM) and 0.14 ± 0.023 µg/ml (0.96 ± 0.16 nM) (Figure 5E). W1F10 IgG1, which preferentially recognizes β1,3-GlcNAc > β1,4-GlcNAc, induced substantially lower levels of phagocytosis at relatively high mAb concentrations (Figure 4D,E and S5) in line with complement deposition results.

### Versatility of the pipeline: identifying functional mAbs against *S. pyogenes*’ Group A carbohydrate

To demonstrate the versatility and broad application of this method, the same experimental setup was used for *S. pyogenes*, an opportunistic pathogen of high priority status for vaccine development, causing nearly half a million deaths globally yearly (18, 19). All *S. pyogenes* strains express a surface glycopolymer, the Group A Carbohydrate (GAC) Lancefield antigen, which consists of repeating subunits of [→3)-α-l-Rha-(1–2)-[β-d-GlcNAc-(1–3)]-α-l-Rha-(1→] of which ∼25% of GlcNAc sidechains are capped with negatively-charged glycerol phosphates (17). The GAC has been instrumental in streptococcal classification and rapid-test diagnostics. Moreover, two different structural variants, i.e. the repeating unit and the PR backbone, are included in preclinical *S. pyogenes* vaccines (20, 21). No human mAbs against these vaccine antigens are currently available as reference in clinical development assays such as opsonophagocytic killing or antibody titer assays. Both GAC variants were chemically synthesized (Figure 6A) and used to identify and sort sGAC-reactive B cells from HD buffy coat. Probe functionality and HD plasma IgG reactivities were successfully verified (Figure 6B and S6). Probe-labeled memory B cells were single cell sorted (0.01%; Figure S7) and after scBCRseq and cloning (17 of 91 [18.7%] cells gave VH-VL pairs), 14 unique mAbs were produced with an IgG1 background and tested for sGAC reactivity. Index sorting of all VH-VL pair matching clones showed memory B cell phenotypes for the sGAC-reactive clones (Figure S8). Seven of these candidates (50%) bound bead-immobilized sGAC (Figure 6C), with 4 clones showing exclusive specificity for sGAC-GlcNAc and not for undecorated PR. The mAbs required the context of the PR backbone for binding since the mAb did not bind to β-1,3-GlcNAc attached to RboP on TarP-modified sWTA (Figure 6C). In addition, 3 clones were reactive to the PR backbone and the reactivity was completely blocked by the modification with GlcNAc (Figure 6C). For further analysis, clones G1C4 and G1E8 were selected as representatives for the GAC-GlcNAc and GAC-PR specificities, respectively.

**Figure 6:**
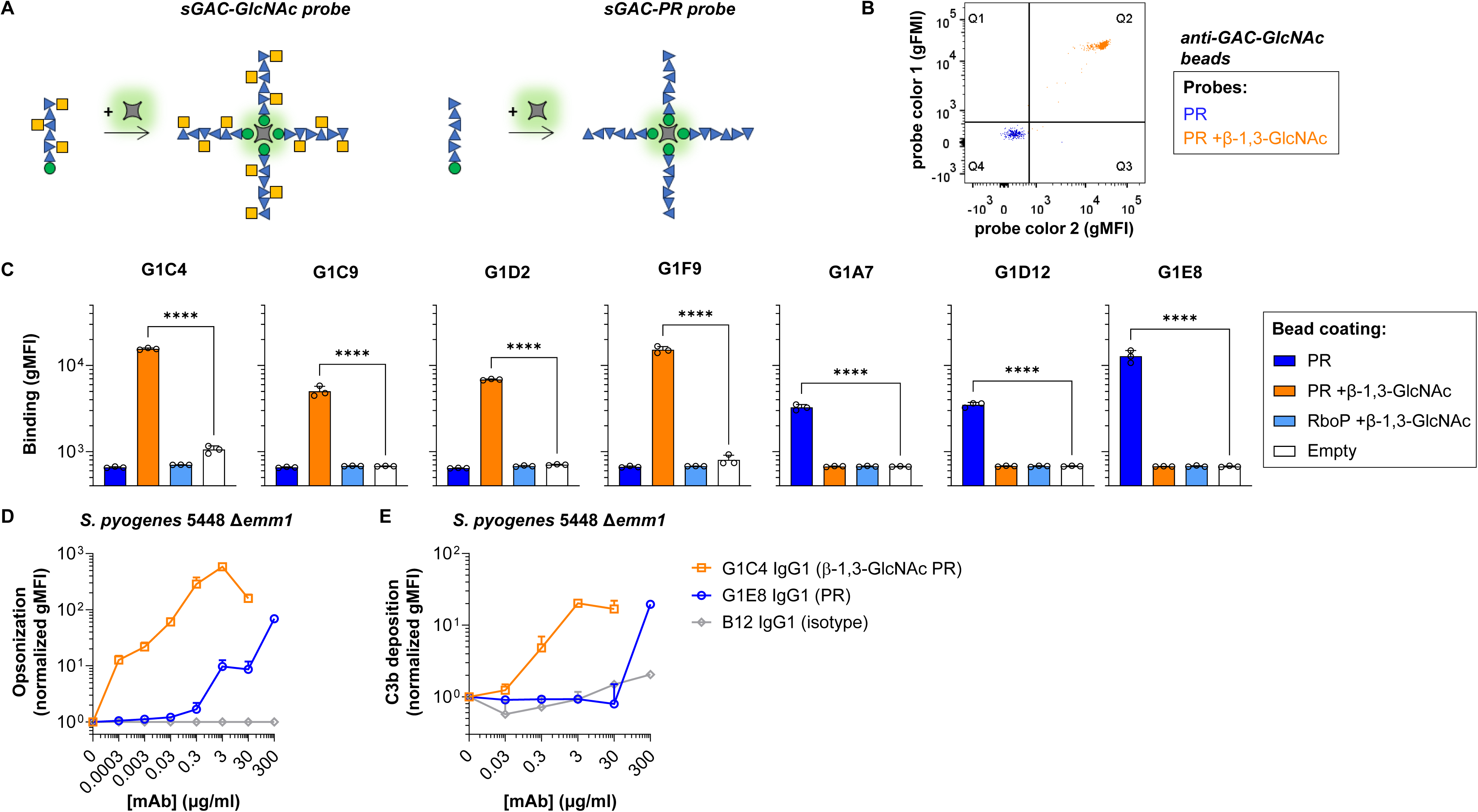
Discovery and characterization of GAC-specific mAbs. **A** Schematic representation of sGAC probe generation for two glycoforms, i.e. PR and PR + β-1,3-GlcNAc. All sGAC probes were made for detection in two different fluorescence channels using streptavidin conjugated to Pe-Cy7 or BV421. **B** sGAC probe binding to protein beads immobilized with goat polyclonal anti-GAC GlcNAc (Ab9191). N.B. no bead coat option was available to test PR specificity. Data in dot plots represent fluorescence signals (fluorophores: PE-Cy7 and BV421) on the beads. Q2 and Q4 comprise, respectively, double positive (dual sGAC probe labeling) and double negative (no sGAC probe binding) beads. Signals within Q1 and Q3 represent aspecific binding of, respectively, Pe-Cy7 and BV421 streptavidin to beads. Histograms are included on the sides of the dot plots to visualize relative amounts of different bead populations within a fluorescent channel. **C** Specificity verification of sGAC-reactive mAbs at equimolar level. Clones were produced in HEK293 Freestyle cells and purified through protein G agarose. Relative binding capacities to sGAC beads were determined at a concentration of 3 μg/ml. Beads coated with RboP +β-1,3-GlcNAc and empty beads were used as controls for cross-reactivity and background, respectively. IgG1 binding to sGAC beads was measured by flow cytometry and data represent the mean gMFI ± s.d. of three independent experiments. One-way ANOVA with Dunnett’s multiple comparisons test was performed to determine significant binding of sGAC-reactive clones to glycan-coated beads compared to empty beads. **** p < 0.0001. Index sort data of all sorted clones can be found in Figure S8. **D** Binding of sGAC-reactive mAbs to natural GAC on *S. pyogenes*. For each glycoform specificity, one mAb clone was selected that showed evident binding to sGAC beads (panel C) and tested for bacterial opsonization. Titration of G1E8 (anti-PR), G1C4 (anti-β-1,3-GlcNAc PR), and B12 (isotype control) to *S. pyogenes* 5448 Δ*emm1*. Bacterial opsonization was determined by measuring IgG1 binding to bacteria using flow cytometry and data represent normalized mean gMFI + s.d. (isotype signals set to 1) of three independent experiments. **E** Effector functions of anti-GAC clones towards *S. pyogenes.* C3b deposition by sGAC-reactive mAbs onto *S. pyogenes* 5448 Δ*emm1*. Data represent C3b binding (normalized gMFI + s.d.) of three independent experiments and was measured by flow cytometry. Fluorescent signals are depicted as a fold change relative to the condition without antibodies to compensate for variation in background signals between biological replicates.

When expressed on the bacterial surface, both G1C4 and G1E8 demonstrated binding to *S. pyogenes* 5448 (lacking IgG-Fc binding M1 protein [Δ*emm1*]) expressing wild-type GAC (AUC 11,188 and 1,044, respectively; Figure 6D). This indicates that both mAbs recognize the more complex and heterogeneous GAC structures on the bacterial surface via the GlcNAc side group (G1C4) or PR (G1E8). G1C4 did not bind to *S. aureus* N315 (AUC 34), which expresses RboP +β-1,3-GlcNAc WTA (Figure S9), confirming the dependence of this anti-GlcNAc mAb on the PR backbone structure. The observation that anti-PR clone G1E8 was able to bind to *S. pyogenes* suggests that on naturally expressed GAC, the decoration with GlcNAc is likely not complete (Figure 6C), i.e. in contrast to the used synthetic fragments not every other rhamnose is modified with GlcNAc.

To test both GAC clones functionally, we studied C3b deposition onto pre-opsonized *S. pyogenes* 5448. Data demonstrate mAb-dependent complement deposition of *S. pyogenes* by G1C4 IgG1 (AUC 483; Figure 6E). The anti-PR G1E8 IgG1 also triggered C3b deposition (AUC 225) but required relatively high mAb concentrations correlating with the lower opsonic capacity of this clone. In conclusion, in line with results for *S. aureus* WTA, synthetic mimics of GAC structures facilitated the discovery of functional mAbs against natural GAC on the surface of *S. pyogenes*.

## DISCUSSION

This study describes an efficient and modular platform to discover human mAbs against bacterial glycans using a workflow (Figure 7) that offers advantages over previous approaches to identify such mAbs. The use of chemically synthesized glycans that allowed us to transition to directly single cell sorting of antigen-specific B cells based on the specificity of interest is critical in this workflow. Our approach eliminates the need for extensive and labor-intensive culturing and specificity verification of unspecified viable B cells, as in previous studies (8, 15). In addition, the synthetic structures are, unlike their natural equivalents, well-characterized and more homogeneous, which allows higher resolution B cell characterization at the epitope level. Furthermore, because of improved stability and homogeneity, synthetic mimics provide a solution for probing unstable target antigens and can deconvolute heterogeneous natural antigenic specificities. This thereby expands the pool of discovered reactivities for less prevailing/dominant or masked epitopes. Interestingly, since WTA-and GAC-reactive antibodies are abundantly present in the circulation of healthy individuals, we could use peripheral B cells from (commercially) available HD buffy coats as input, instead of limited amounts of convalescent blood from patients. In case of rare reactivities, HD input can be upscaled by sampling more residual blood collection material, something that is virtually impossible when relying on patient material. However, when the antibodies of interest only arise after infection, patient material could also be used as input for the pipeline. Finally, compared to other conventional mAb discovery strategies, e.g. hybridoma and llama immunization, this setup using human material promotes the transition to animal-free research.

**Figure 7:**
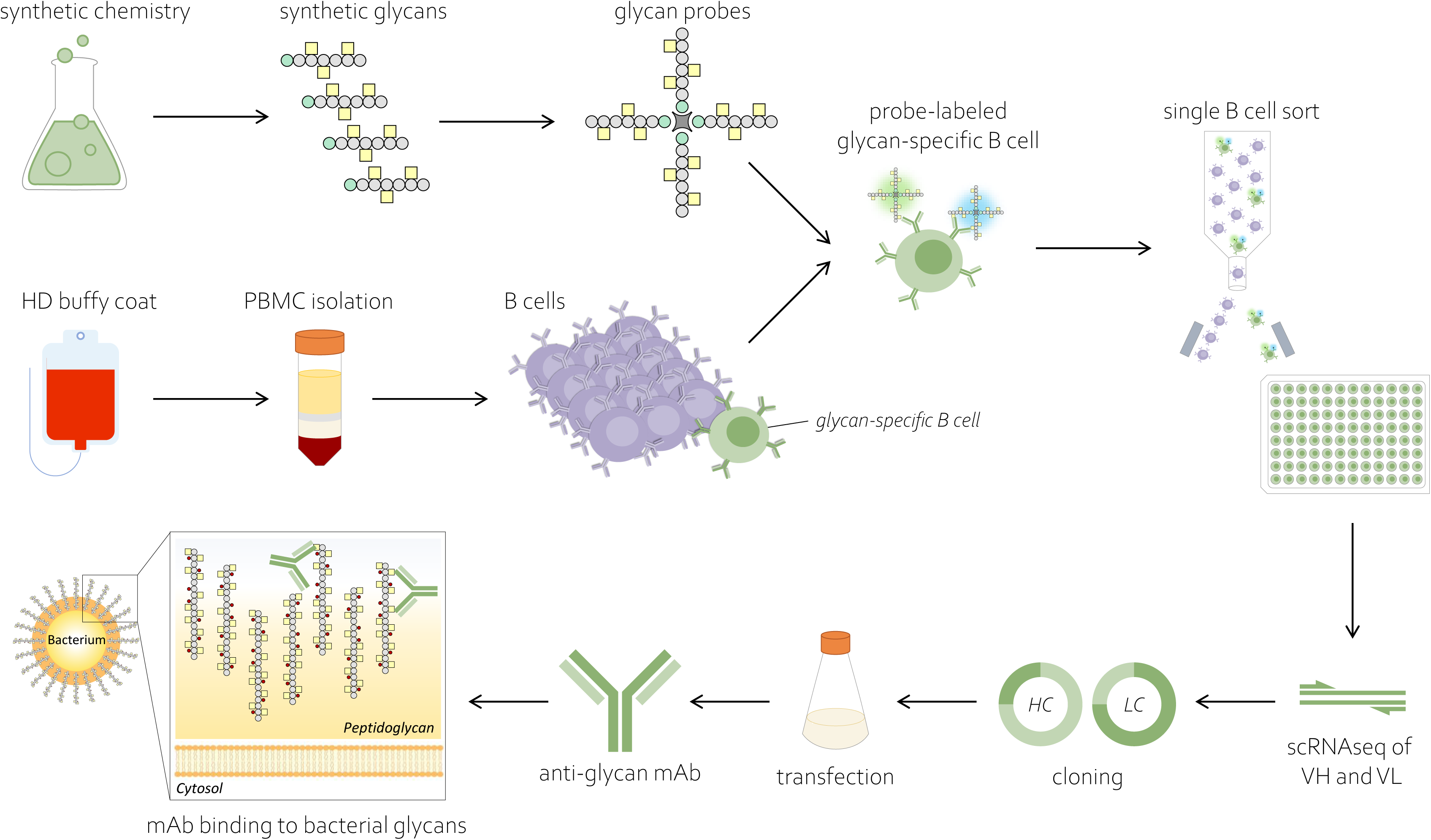
Schematic overview of experimental workflow for identifying glycan-specific antibodies against bacterial pathogens using synthetic glycan fragments.

Using our workflow, we successfully identified mAbs that target surface glycans of two different priority pathogens; 16 anti-WTA clones (10 unique) from 78 sorted B cells, and 7 unique anti-GAC clones from 91. For *S. aureus*, the anti-WTA clones recognized GlcNAc attached through the three different conformations, i.e. β-1,4-, β-1,3-, and α-1,4-GlcNAc. Notably, no clones were identified that recognized WTA GlcNAc universally regardless of conformation indicating differing immunorecognition of α- and β-GlcNAc. The specificities found here are similar to those of previously identified human anti-WTA clones that exclusively target WTA via GlcNAc (8, 15) and corroborates earlier work that GlcNAc is the dominant epitope in WTA-specific antibody responses (22). The discovered clones appear to bind specifically to *S. aureus* WTA GlcNAc since interactions required the context of the RboP backbone. To our knowledge, the W1F10 clone exhibits unique epitope recognition by discriminating β-1,3-GlcNAc from β-1,4-GlcNAc. Previously, another anti-β-GlcNAc clone (Genentech clone 6292 (15)) showed similar distinctive binding characteristics, but in this case with a preference for β-1,4-GlcNAc over β-1,3-GlcNAc (10).

Several monoclonal anti-GAC-GlcNAc formulations are commercially available, but all are sourced from immunized animals rendering them immunogenic when applied in human settings. The anti-GAC mAbs in this study are of human origin and therefore more compatible with *in vivo* diagnostic, therapeutic and prophylactic use in humans. Recently published preliminary work has also identified human anti-GAC mAbs by single cell sorting and sequencing of B cells from patient tonsillectomy material through staining with natural GAC structures isolated from *S. pyogenes* (23). In this study, the identified clones were reactive against the GlcNAc epitope but mainly after increasing binding avidity and predominantly irrespective of the GAC context (i.e. PR backbone). The specificities of our mAbs for GAC-GlcNAc depend on GlcNAc interaction next to PR contacts, as no cross-reactivity to RboP-attached GlcNAc residues could be observed. In addition, our clones did not require avidity enhancement for *S. pyogenes* binding and are sourced from one healthy donor instead of patient material. Moreover, synthetic GAC allowed for epitope-specific B cell staining, yielding dominant GlcNAc reactivities as well as less dominant epitopes, i.e. PR. This exemplifies the additional value of using well-defined homogenic synthetic mimics over natural equivalents, especially to find specificities of interest that are naturally overshadowed by dominant epitopes.

In the past, antibody reactivity against the GAC-GlcNAc epitope has been linked to the pathogenesis of rheumatic heart disease due to the proposed cross-reactivity to human GlcNAc expressed on cardiac and neuronal tissue (24–27). Alternatively, recognition of GAC via alternative epitopes, e.g. PR, may provide safer targeting strategies in clinical settings, because PR is non-mammalian. G1E8 can be considered for this as this clone targets GAC via PR (a unique specificity that has never been described before) and importantly is able to recognize *S. pyogenes* expressing PR with GlcNAc modifications. However, compared to anti-GAC GlcNAc clone G1C4, opsonic saturation levels are lower for G1E8 reflecting PR’s lower availability and/or lower mAb affinity towards the epitope. Nonetheless, G1E8 was able to induce C3b deposition onto *S. pyogenes* though at relatively high mAb concentrations. To optimize G1E8 functionality, crafting of the VH/VL domains and (glyco)engineering of the IgG-Fc tail may be employed to improve affinity, avidity and/or effector functions.

The general consensus on the GAC structure is that GlcNAc is present on every 1,3-rhamnose-1,2-rhamnose repeating unit (28). This is also the structure that was mimicked with the sGAC fragment. As we observed for clone G1E8, recognition of the PR backbone was blocked upon saturated GlcNAc decoration. In contrast, G1E8 was able to opsonize *S. pyogenes* expressing wild-type GAC. This suggests that decoration of PR structures on *S. pyogenes* may not always be as dense as dogmatized in literature. Alternatively, PR is also present on the *S. pyogenes* surface alongside GlcNAcylated-GAC, which is possible since GlcNAc attachment occurs extracellularly (28). Either way, these data suggest that PR is an available and therefore relevant target epitope for antibodies and mAbs.

We used a cocktail of sWTA glycoforms to identify as many different antibody specificities as possible. Since these glycoform probes were generated with identical fluorophore combinations, the B cells could only be specified on antigen but not epitope level prior to sorting. Alternatively, sWTA glycoforms can also be probed individually or labeled with unique fluorophore pairs to specify B cells with a higher resolution at epitope level. Especially for the identification of more rare specificities, exclusion of immunodominant epitopes that potentially compete for probe binding would be desirable. Complementary, undesired glycoform specificities can also be excluded from the sort when probes are used for depletion via dump gating.

We expressed the VH and VL domains of the newly identified clones as mAbs of the IgG1 subclass, which may be different from their original isotype or subclass. We chose IgG1 since this subclass efficiently elicits Fc-mediated effector functions such as complement activation and IgG-receptor mediated uptake by neutrophils. Indeed, all mAbs triggered Fc-mediated functionalities and thereby demonstrated their prophylactic and therapeutic potential, which can be further optimized by applying (glyco)engineering strategies (29). Potentially, the clones could also be useful diagnostically in mAb-based point of care tests, *in vivo* imaging, and in characterization of the infecting pathogen with high resolution, i.e. at epitope level. For this, effector functions should be avoided and thus require the clones to be engineered in an inert/less potent antibody framework, e.g. Fc-silent, F(ab’)_2_/Fab fragments (29).

Although our workflow effectively isolated uncommon glycan-specific clones from a large, heterogeneous B cell pool, the yield can be further optimized. First, unreactive or very low-affinity B cell clones can be excluded from the B cell sorting process by adjusting the gating strategy, ensuring that the sorted dual-labeled B cells originate exclusively from the memory compartment. Second, the lack of matching VH or VL sequences to form complete VH–VL pairs was a considerable limitation to our mAb discovery and could potentially be optimized by incorporating additional (newly identified) germline sequences to the already well-established primer panels used for nested PCR. Third, the number of reactive mAbs could be increased by expressing each clone in the isotype and subclass corresponding to its original B cell phenotype. This may uncover antigen specificity that is not apparent in the IgG1 format, for example due to enhanced avidity (e.g. IgM) or increased structural flexibility (e.g. longer hinge regions).

In conclusion, this study exemplifies the use of synthetic glycopolymers as powerful tools to facilitate the discovery of human mAbs against bacterial glycans with high resolution. The setup we describe provides an efficient and modular workflow to generate anti-glycan mAbs from HD human B cells. In this way, we identified anti-WTA and anti-GAC mAbs reactive against varying (unique) epitopes on the respective antigens. Taken together, our work paves the way for the discovery of additional glycan-specific mAbs, which can contribute to improved and tailored diagnostics, prevention, and therapeutic applications for *S. aureus*, *S. pyogenes* and other (AMR) pathogens.

## MATERIALS AND METHODS

### Bacterial Strains and Culture Conditions

*S. aureus* strains (Table 2) were grown overnight with agitation in Tryptic Soy Broth (TSB). *S. pyogenes* (Table 2) were grown overnight without agitation in Todd-Hewitt broth with yeast (THY). The next day, overnight cultures were subcultured in fresh medium, grown to mid-exponential phase (optical density at 600 nm [OD_600_) of 0.6], centrifuged, and resuspended in PBS at an OD_600_ of 0.4.

### Human Material

Buffy coats were ordered from Sanquin Blood Supply (Amsterdam, the Netherlands). Upon delivery, PBMCs were isolated through Ficoll-Paque Plus density gradient centrifugation (30). The peripheral blood mononuclear cell (PBMC) fractions were stored in freeze medium at -80°C until further use. Human neutrophils were isolated from heparinized blood drawn from healthy volunteers (bloedafnames ten behoeve van biomedische onderzoekslaboratoria [BACON] protocol) by Ficoll-Histopaque density gradient centrifugation (31).

### Synthetic Glycan Production, Enzymatic Glycosylation and Bead Coupling

The biotinylated sWTA hexameric structures with a poly-RboP backbone structure were generated through synthetic chemistry as previously described by Ali et al. (32). The biotinylated linear hexamers with a backbone consisting of repeating rhamnose units and GlcNAc side groups, resembling the *S. pyogenes* GAC, were generated in a similar manner (33). Enzymatic glycosylation and bead coupling of sWTA were performed as previously described by van Dalen et al. (10). In short, 0.17 mM unmodified RboP hexamers were incubated with recombinant TarS, TarM, or TarP enzyme (6.3 µg/ml) and UDP-GlcNAc substrate (2 mM) in glycosylation buffer (containing 15 mM HEPES, 20 mM NaCl, 1 mM EGTA, 0.02% Tween-20, 10 mM MgCl_2_, 0.1% BSA, pH 7.4) for 2 hours at room temperature. After incubation, glycosylated structures were heat-inactivated at 65°C for 20 minutes to abolish enzymatic activity. Next, the biotinylated sWTA/sGAC structures were incubated with streptavidin-coated magnetic Dynabeads (M280; Thermo Scientific) for 15 minutes at room temperature with shaking (600 rpm). After incubation, beads were washed three times with PBS using an Eppendorf magnet and eventually resuspended in PBS supplemented with 0.05% Tween-20 and 0.1% bovine serum albumin (BSA), and stored at 4°C until further use.

### Generation and Testing of sWTA/sGAC Probes

Biotinylated sWTA or sGAC were incubated with streptavidin conjugates (BB515 and AF647 or BV421 and Pe-Cy7, respectively; Thermo Scientific) in a molar ratio of 4:1 for at least 90 minutes on ice in the dark, resulting in fluorescent probes with a concentration of 450 nM. Subsequently, a surplus of free biotin (0.1 mM) was added to the probes to saturate potential unoccupied binding sites on streptavidin. Probes were stored at 4°C until further use.

### B Cell Preparation and Single Cell Sorting

The B cell sorting was based on the setup as previously described in Brouwer et al.(16) with some adaptations. The PBMC fraction was enriched for B cells using the Pan B Cell Isolation Kit, human (130-101-638, Miltenyi Biotec), according to manufactureŕs instructions. After Pan B cell negative isolation, B cells were washed and resuspended in buffer (PBS supplemented with 2% fetal calf serum and 1 mM EDTA) to a concentration of 1.2 × 10^8^ cells/ml. Subsequently, cell suspension was diluted (1:1, v/v) with fluorescent sWTA/sGAC probes and incubated in the dark for 1 hour on ice. Cells were washed, resuspended in antibody mixture (Table 1) to a concentration of 1 × 10^8^ cells/ml, and incubated in the dark for 30 minutes on ice. Cells were washed and resuspended in buffer to a final concentration of 20 × 10^6^ cells/ml. Flow cytometry and single B cell sorting was performed on a FACSAria Illu Sorp flow cytometer (BD Biosciences). The used gating strategy resulted in sorting of single CD19+ dump-B cells double positive for both probe labels, i.e. BB5151 and AF647 for sWTA, and BV421 and Pe-Cy7 for sGAC (dump gate encompassed live/dead marker and non-B cell markers: CD3, CD4, CD14 and CD16). Dual positivity for the probe reactivity was used to minimize the amount of false-positive B cells specific for one of the two fluorochromes instead of WTA/GAC. For further characterization, cells were also labeled for B cell maturation markers IgD and CD27. Gated cells were single cell sorted into a 96-wells PCR plate and stored at -80°C until further use.

**Table 1:** Details of reagents used in antibody mixture for B cell labeling. AF, Alexa Fluor. eF, eFluor. APC, Allophycocyanin. BV, Brilliant Violet. PE, Phycoerythrin

| Product name | Conjugate | Clone | Reference | Company | Dilution used |
| --- | --- | --- | --- | --- | --- |
| anti-human CD19 Antibody | AF700 | HIB19 | 302226 | Biolegend | 1:8 |
| Fixable Viability Dye | eF780 | n.a. | 65-0865-14 | eBioscience | 1:1000 |
| CD4 Monoclonal Antibody | APC eF780 | OKT4 | 47-0048-42 | eBioscience | 1:33 |
| CD3 Monoclonal Antibody | APC eF780 | UCHT1 | 47-0038-42 | eBioscience | 1:33 |
| CD14 Monoclonal Antibody | APC eF780 | C1D3 | 47-0149-42 | eBioscience | 1:20 |
| CD16 Monoclonal Antibody | APC eF780 | CB16 | 47-0168-42 | eBioscience | 1:50 |
| anti-human IgD Antibody | BV785 | IA6-2 | 348242 | Biolegend | 1:5.5 |
| anti-human CD27 Antibody | PE | O323 | 302807 | Biolegend | 1:5.5 |

*After diluting the abovementioned reagents in PBS supplemented with 2% fetal calf serum and 1mM EDTA, Brilliant Stain Buffer was added 1:8 for proper BV staining*

### Single Cell B Cell Receptor Sequencing

After the plate with single-sorted cells was thawed on ice, B cells were lysed by adding 20 μl of lysis buffer (containing 20 units RNAse inhibitor, First Strand Superscript III buffer and 6.25 mM DTT, Invitrogen) to each well, and the plate was incubated at -80°C for 24 hours. To generate cDNA from the B cell RNA, a reverse transcriptase (RT)-PCR was performed, as described previously (16, 30). To each well, 6 μl of RT-mix (200 ng random hexamer primers, 2 mM dNTP, 50 units SuperScript III RTase, Invitrogen) was added and subjected to PCR with the following settings: 10 minutes at 42°C; 10 minutes at 25°C; 1 hour at 50°C; 5 minutes at 95°C; infinite 8°C.

Three rounds of successive nested PCR reactions were performed to amplify specifically the BCR VH and VL sequences from the cDNA, as described previously (16, 30). Restriction sites flanking the VH or VL sequences were incorporated during the last round of PCR using primers. After PCR clean-up, the VH (AgeI × SalI) and VL (AgeI × BsiWI or XhoI) PCR products were cloned into mammalian expression plasmids encoding the human IgG1 constant domains CH1-3 and the kappa light chain, respectively, through double restriction and ligation. Sanger sequencing with BDT1.1 was performed on the generated constructs to identify the following nucleotide sequence of the VH and VL domains

### Recombinant Expression of Monoclonal Antibodies

The newly identified WTA/GAC-specific BCR clones, Genentech clones 4497 and 4461 (patent WO2014193722A1) and isotype control (anti-HIV-1 gp120, clone B12) were recombinantly produced as human IgG1, as described previously by Aartse et al. (34), with some adaptations. For pilot-scale productions, expression plasmids encoding the VH+IgG1 and VL+kappa/lambda light chain sequences were 1:1 w/w co-transfected into HEK293T cells and cell supernatants were harvested 3 days post-transfection. IgG1 production levels of were determined using protein A beads assay. For larger scale productions, expression plasmids encoding the heavy and light chain sequences were 1:1 w/w co-transfected into HEK293 Freestyle cells using PEImax. The mAbs were harvested through centrifugation of the cell suspension after 5 days shaking at 37°C and 8% CO_2_. The supernatant was filtered through a 0.2 um Steritop bottle top filter (Millipore), protein G agarose beads (Pierce) were added, and incubated overnight rolling at 4°C. The next day the supernatant was applied to a column where the beads were allowed to sediment and being washed with PBS. The mAbs were eluted from the protein G agarose beads with 0.1 M Glycine pH 2.5 and neutralized with 1 M Tris pH 8.7 (9:1, v/v). The eluted fraction was subsequently buffer-exchanged to PBS using a Vivaspin concentration tube with 100K MWCO (Sartorius). The mAbs were stored at 4°C until further use.

### Bead Assays

The mAb specificities were verified using a bead assay where magnetic beads (1 × 10⁵) coated with (un)modified sWTA/sGAC hexamers were incubated with the mAbs serially diluted in PBS supplemented with 0.1% BSA and 0.05% Tween-20. After 20 minutes shaking (600 rpm) at 4°C, beads were pelleted using a plate magnet and the antibody mixture was aspirated. Subsequently, beads were incubated with 1:1,000 diluted AlexaFluor488-conjugated protein G (P11065, Thermo Scientific) or 1:500 PE-conjugated mouse anti-Human IgG1 Fc (HP6001; Southern Biotech). After 20 minutes in the dark and shaking (600 rpm) at 4°C, the beads were pelleted and supernatant was aspirated. Finally, the beads were resuspended in PBS supplemented with 0.1% BSA and 0.05% Tween-20 and measured in a FACSCanto flow cytometer (BD). Fluorescent signals are depicted as gMFI values or as a fold change relative to the condition without antibodies: gMFI mAb ÷ gMFI no mAb, to compensate for technical variation.

Probe functionalities were tested using beads coated with antibodies against WTA or GAC. For WTA, 1 × 10⁵ protein A-coated magnetic Dynabeads (10001D; Thermo Scientific) were incubated for 20 minutes shaking (600 rpm) at 4°C with 20 µg/ml human IgG1-based mAbs, i.e. 4497 or 4461 human IgG1 to verify respectively β-GlcNAc and α-GlcNAc specificities. Isotype IgG1 (B12) was included for negative control purposes. For GAC, protein A beads were first coated with Rabbit anti-Goat IgG (H+L) Secondary Antibodies (31105, Thermo Scientific) and subsequently with 30 µg/ml goat polyclonal *S. pyogenes* Group A carbohydrate antibody (ab9191; Abcam) as protein A has negligible affinity towards goat IgG-Fc. After coating with antibodies, beads were washed and incubated glycoform-specifically with fluorescent sWTA or sGAC probe pairs (different conjugates at 1:1 ratio) diluted in PBS supplemented with 0.1% BSA and 0.05% Tween-20. After 20 minutes in the dark and shaking (600 rpm) at 4°C, the beads were pelleted and supernatant was aspirated. Finally, the beads were resuspended in PBS supplemented with 0.1% BSA and 0.05% Tween-20.

IgG1 levels of pilot-scale productions were determined by incubating serial dilutions of HEK293T cell supernatants 1:1 (v/v in PBS supplemented with 0.1% BSA and 0.05% Tween-20) with 1 × 10⁵ protein A-coated magnetic Dynabeads (10001D; Thermo Scientific). After 20 minutes shaking (600 rpm) at 4°C, beads were pelleted using a plate magnet and the antibody mixture was aspirated. Subsequently, beads were incubated with 1:1,000 diluted AlexaFluor488-conjugated protein G (P11065, Thermo Scientific). After 20 minutes in the dark and shaking (600 rpm) at 4°C, the beads were pelleted and supernatant was aspirated. Finally, the beads were resuspended in PBS supplemented with 0.1% BSA and 0.05% Tween-20 and measured in a FACSCanto flow cytometer (BD).

### Bacterial Binding Assay

To assess binding to native WTA/GAC expressed on bacterial surfaces, mAbs were serially diluted in reaction buffer (PBS supplemented with 0.1% BSA) and incubated (1:1, v/v) with bacteria (OD_600_ = 0.4 in PBS) to a final volume of 25 μl. After incubating for 20 minutes on a rotating platform (600 rpm) at 37°C, the 96 wells plate was washed by adding 100 μl reaction buffer to each well and centrifugation (2,684 rcf, 10 minutes), the supernatant aspirated, and 25 μl AlexaFluor488-conjugated goat anti-human IgG F(ab)_2_ fragments (2042-30; Southern Biotech) 1:200 diluted in PBS 0.1% BSA were added. After another incubation step of 20 minutes on a rotating platform at 37°C in the dark, the plate was washed, the supernatant aspirated, and bacteria were fixed by resuspending in 150 μl filter-sterile PBS supplemented with 1% paraformaldehyde (Sigma-Aldrich). Samples were measured in a FACSCanto flow cytometer (BD Biosciences). IgG1 isotype control (B12) was included to define geometric Mean Fluorescence Intensity (gMFI) values generated by residual aspecific binding of IgG1 (e.g. in N315 strains where only the *spa* gene is knocked out, and not the *sbi* gene encoding IgG1 binding protein Sbi). Data were normalized by setting IgG1 isotype signals to 1 to remove aspecific binding signals.

### Complement Deposition Assay

The pre-made aliquots of bacterial cultures of OD_600_ = 0.4 (Table 2) were used for the C3b complement deposition assays. IgG/IgM depleted pooled human serum (34010 Pel-Freez Biologicals), stored at - 70°C was used as a complement source. The bacterial suspension and mAbs (10-fold serial dilution from 30 or 300 to 0 µg/ml final concentration) were mixed at a 1:1 ratio (end volume 25 µL) in a sterile round bottom 96-well plate and incubated for 20 min at 4°C. After incubation, the wells were washed with 125 µL of Roswell Park Memorial Institute medium (RPMI) + 0.1 % BSA (Serva) and centrifuged (4,000 rpm) for 10 min at 4°C. The supernatant was carefully aspirated, and the pellet was re-suspended in 50 µL of 1% IgG/IgM-depleted human serum. The plates were incubated spinning (525 rpm) for 30 min at 4°C. After the incubation, the plates were washed with 125 µL of RPMI + 0.1 % BSA and centrifuged (3,500 rpm) at 4°C for 10 min. The supernatant was carefully aspirated, and the pellet was re-suspended in 25 µL of FITC F(ab’)2 Goat Anti-Human Complement C3 (Mpbio) diluted 1:50 in RPMI + 0.1 % BSA. Subsequently, the plates were incubated for 20 min at 4°C in the dark and the washing step was repeated. Finally, the pellet was fixed with 150 µL of 1% paraformaldehyde (Sigma-Aldrich), diluted in RPMI + 0.1 % BSA. The plates were stored at 4°C in the dark until flow cytometer measurements were conducted. The flow cytometer measurements were performed with the BD FACS CANTO II (BD Biosciences). The data was collected with the BD FACS Diva Software (BD Biosciences) and analyzed with FlowJo 10, FCSexpress 7 (DeNovo Software). Fluorescent signals are depicted as a fold change relative to the condition without antibodies: gMFI mAb ÷ gMFI no mAb, to compensate for variation in background signals between biological replicates.

**Table 2:**
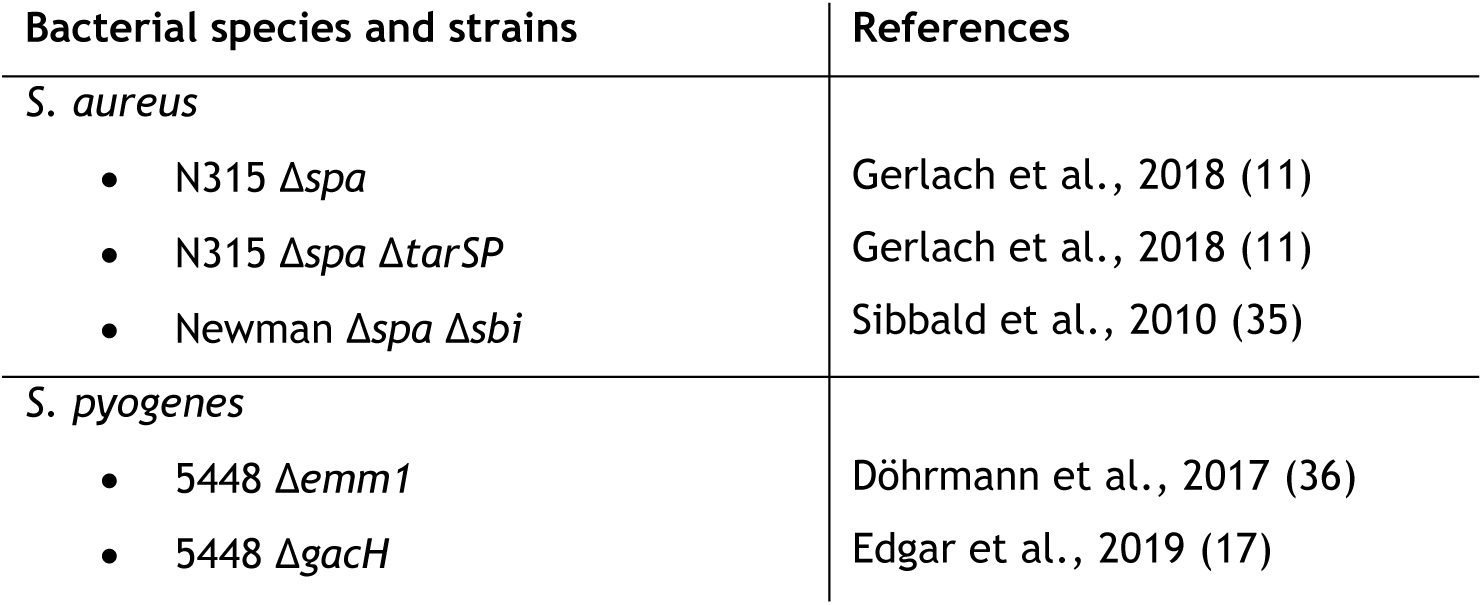
List of the bacterial species and strains used in this study.

| Bacterial species and strains | References |
| --- | --- |
| *S. aureus* |  |
| • N315 $\Delta spa$ | Gerlach et al., 2018 (11) |
| • N315 $\Delta spa \Delta tarSP$ | Gerlach et al., 2018 (11) |
| • Newman $\Delta spa \Delta sbi$ | Sibbald et al., 2010 (35) |
| *S. pyogenes* |  |
| • 5448 $\Delta emm1$ | Döhrmann et al., 2017 (36) |
| • 5448 $\Delta gacH$ | Edgar et al., 2019 (17) |

### Phagocytosis Assay

GFP-expressing *S. aureus* Newman Δ*spa* Δ*sbi* strain was freshly grown to midlog (OD_600_ = 0.6), diluted in RPMI +0.05% HSA (OD_600_ = 0.4) and used for phagocytosis assays. IgG/IgM depleted pooled human serum stored at -70°C was used as a complement source at a concentration of 1%. The bacterial suspension (10x diluted), mAbs, and serum were mixed at a 2:1:1 ratio (end volume 40 µL) in a sterile round bottom 96-well plate and incubated for 15 minutes, at 37°C, shaking (700 RPM, on plate shaker). After incubation, 10 μl of neutrophils were added in an E:T ratio of 1:10 to the wells and again incubated for 15 minutes, at 37°C, shaking. Subsequently, cells were fixed by adding 2% paraformaldehyde in RPMI to the wells in 1:1 ratio. The plates were stored at 4°C in the dark until flow cytometer measurements were conducted. The flow cytometer measurements were performed with the BD FACS CANTO II (BD Biosciences). The data was collected with the BD FACS Diva Software (BD Biosciences) and analyzed with FlowJo 10, FCSexpress 7 (DeNovo Software).

### Data Analyses

FlowJo was used to analyze flow cytometry data, generate dot plots, and export geometric mean fluorescence intensities as Excel tables for further analysis. To analyze each sorted B cell per specific well, index sorting data was exported from the corresponding experiments in FACS Diva as a .csv file with geometric mean values of all relevant parameter channels. The data was consecutively analyzed in Excel. Graphpad Prism was used to generate graphs, determine AUC and IC_50_, and perform curve fitting and statistical tests.

## Acknowledgements

The authors would like to thank volunteers for donating blood; L.H.E. van Huijkelom, S. Bovenkerk-Man, and E.M. Struijf for arranging blood donations and isolating neutrophils; J.L. Snitselaar and M.J. van Breemen for culturing and preparing the HEK293T and Freestyle cells; W. Olijhoek for sharing his practical knowledge on RT and nested PCR reactions; Microscopy and Cytometry Core Facility - location AMC (MCCF-AMC).

## Author Contributions

A.R. Temming conceptualized, designed and executed all experiments reported in this study, and wrote the manuscript. M. Claireaux and G. Kerster prepared PBMCs, facilitated designing and setting up B cell sorting. S.E. Groenewege generated bead assay data for the identified mAbs under supervision of A.R. Temming. T. Voskuilen, Z. Wang and J.D.C. Codée produced and provided synthetic glycans. M.J. van Gils conceptualized and facilitated setting up scBCRseq experiments and mAb production. N.M. van Sorge conceptualized and supervised the study, and wrote this manuscript. The manuscript was approved and critically reviewed by all co-authors.

---

**Figure S1: Profiling healthy donor buffy coat plasma for sWTA-specific antibodies**

Isotype profiling of healthy donor (HD) plasma for sWTA reactivity. Plasma was titrated and incubated with beads coated with RboP +β-1,4-GlcNAc, +α-1,4-GlcNAc, or +β-1,3-GlcNAc. Bead-bound IgG, IgM, and IgA were measured using flow cytometry. Data represent geometric mean fluorescence intensities after subtraction of the gMFI value belonging to 0% dilution to remove background variation (i.e. ΔgMFI = gMFI mAb – gMFI no mAb).

**Figure S2: Gating strategy (A) and maturation status (B) of sorted sWTA probe-labelled B cells**

Cut off values for maturation marker expression are indicated by dotted lines in order to distinguish unswitched (IgD+ CD27+) and switched (IgD-CD27+) memory B cells, and naïve B cells (IgD+ CD27-).

**Figure S3: IgG1 levels in pilot-scale productions**

Human IgG1 levels in HEK293T cell supernatants of 16 B cell-derived mAbs. Supernatant dilutions were incubated with magnetic protein A-coated beads to determine mAb production levels as measured by flow cytometry. Data represent area under the curve (AUC) values of titration curves ranging from 50% to 6.25% supernatant dilutions. W1G5 was the only clone that exhibited negligible production and was therefore excluded from the specificity screening in **main Figure 2**.

**Figure S4: Index sort of sWTA probe-reactive sorted B cells**

sWTA probe labeling profiles (left) and maturation status (right) of all sorted B cells. Cells are categorized based on whether the BCR VH-VL pair was identified (or not) for mAb production. The clones with matching VH-VL pairs were further subdivided into those that could be produced (or not) and their reactivity towards sWTA beads as depicted in **main Figure 2**. Cut off values for maturation marker expression are indicated by dotted lines in order to identify unswitched (IgD+ CD27+) and switched (IgD-CD27+) memory B cells and naïve B cells (IgD+ CD27-).

**Figure S5: Neutrophil-mediated phagocytosis of GFP-expressing *S. aureus* Newman *Δspa Δsbi* by anti-sWTA mAbs**

Each panel represents an individual biological replicate. Reactions were performed in the presence of 1% IgG-IgM-depleted human serum. Displayed data represent percentages of GFP-positive neutrophils. Curves were generated using nonlinear dose-response fitting model and used to determine the IC_50_ values depicted in main Figure 5E. Horizontal dotted lines indicate the y-value of 50% effect ([[curve top − bottom] ÷ 2] + bottom) and the mAb concentration of the interception point (cross) with the curve corresponds to IC_50_ (intercept of vertical dotted line with x-axis).

**Figure S6: sGAC-specific antibody profiling healthy donor plasma**

Isotype profiling of HD plasma for sGAC reactivity. Plasma was titrated and incubated with beads coated with PR or PR +β-1,3-GlcNAc sGAC glycoforms. Bead-bound IgG, IgM, and IgA were measured using flow cytometry. Data represent geometric mean fluorescence intensities after subtraction of the gMFI value belonging to 0% dilution (i.e. ΔgMFI = gMFI mAb – gMFI no mAb) to remove background variation.

**Figure S7: Gating strategy (A) and maturation status (B) of sorted sGAC probe-labelled B cells** Cut off values for maturation marker expression are indicated by dotted lines in order to distinguish unswitched (IgD+ CD27+) and switched (IgD-CD27+) memory B cells.

Fig. S8: **Index sort of sGAC probe-labelled sorted B cells**

sGAC probe labeling profiles (left) and maturation status (right) of all sorted B cells. Cells are categorized based on whether the BCR VH-VL pair was identified (or not) for mAb production. The clones with matching VH-VL pairs were further subdivided into those that could be produced (or not) and their confirmed reactivity towards sGAC beads as depicted in main Figure 6C. Cut off values for maturation marker expression are indicated by dotted lines in order to identify unswitched (IgD+ CD27+) and switched (IgD-CD27+) memory B cells and naïve B cells (IgD+ CD27-).

**Figure S9: sGAC β-1,3-GlcNAc-reactive mAb G1C4 is not cross-reactive to β-1,3-GlcNAc on RboP-based *S. aureus* WTA**

