## Supplementary figures and images for "Platform for Identifying Human Glycan-Specific Antibodies Against Bacterial Pathogens using Synthetic Glycan Fragments"

### Supplemental figures

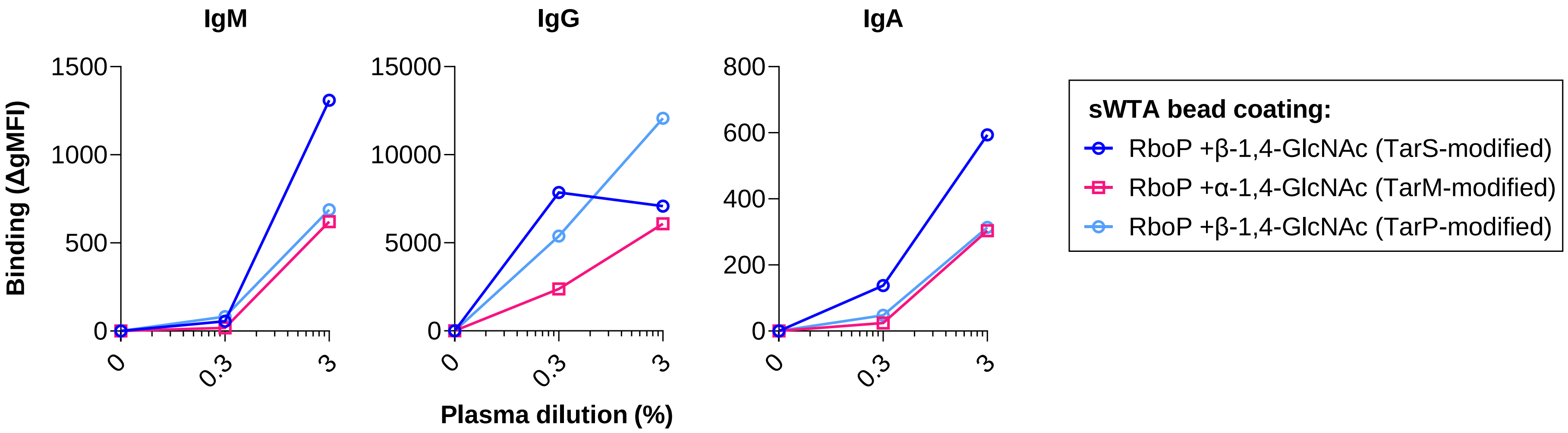

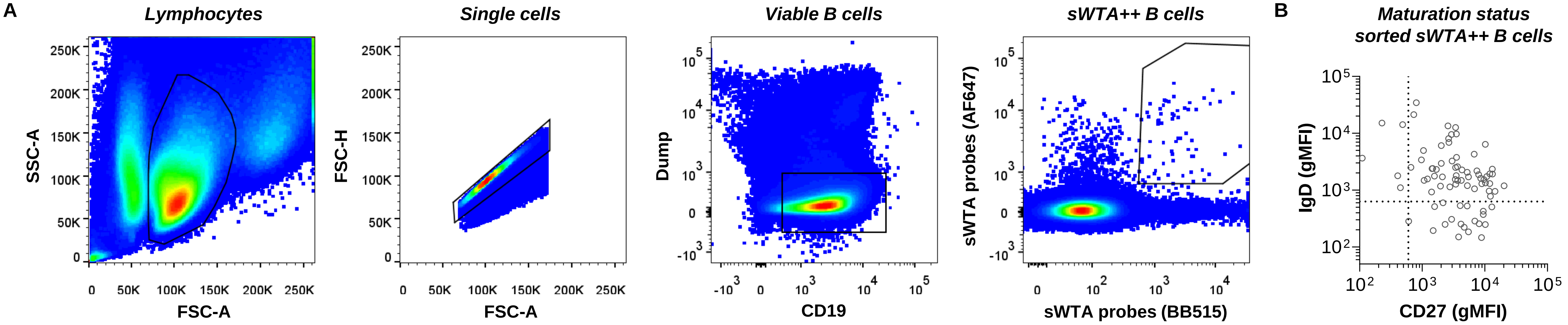

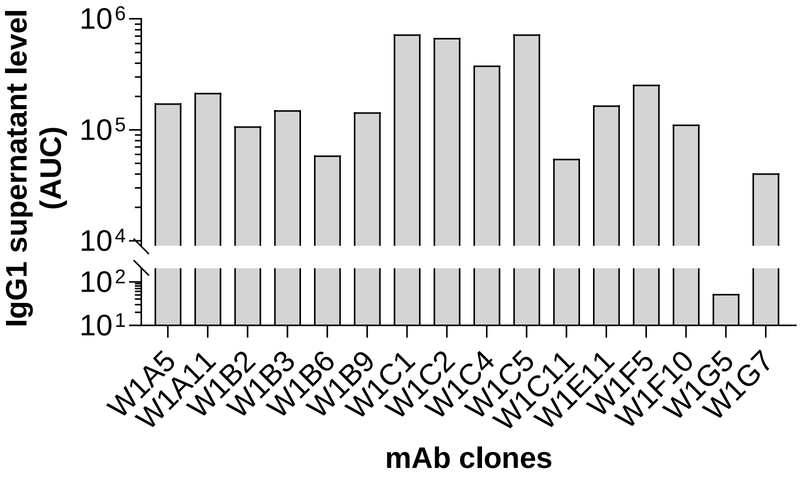

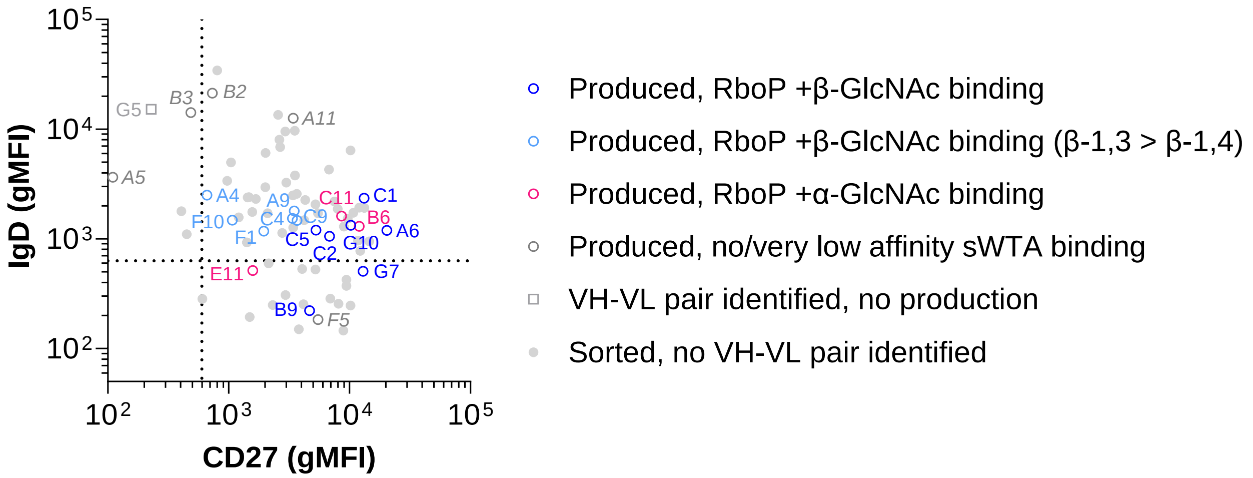

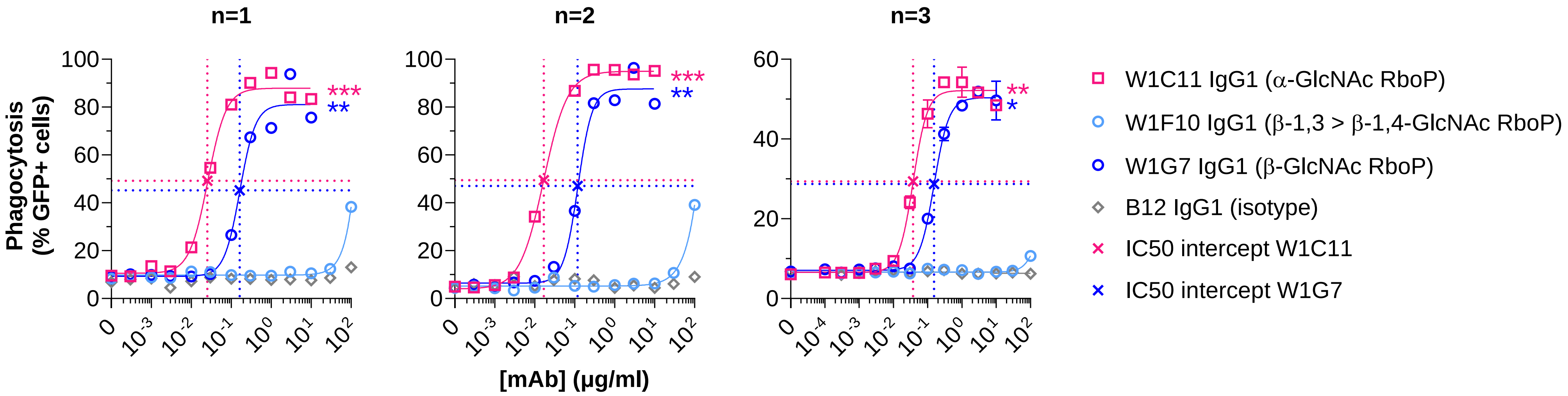

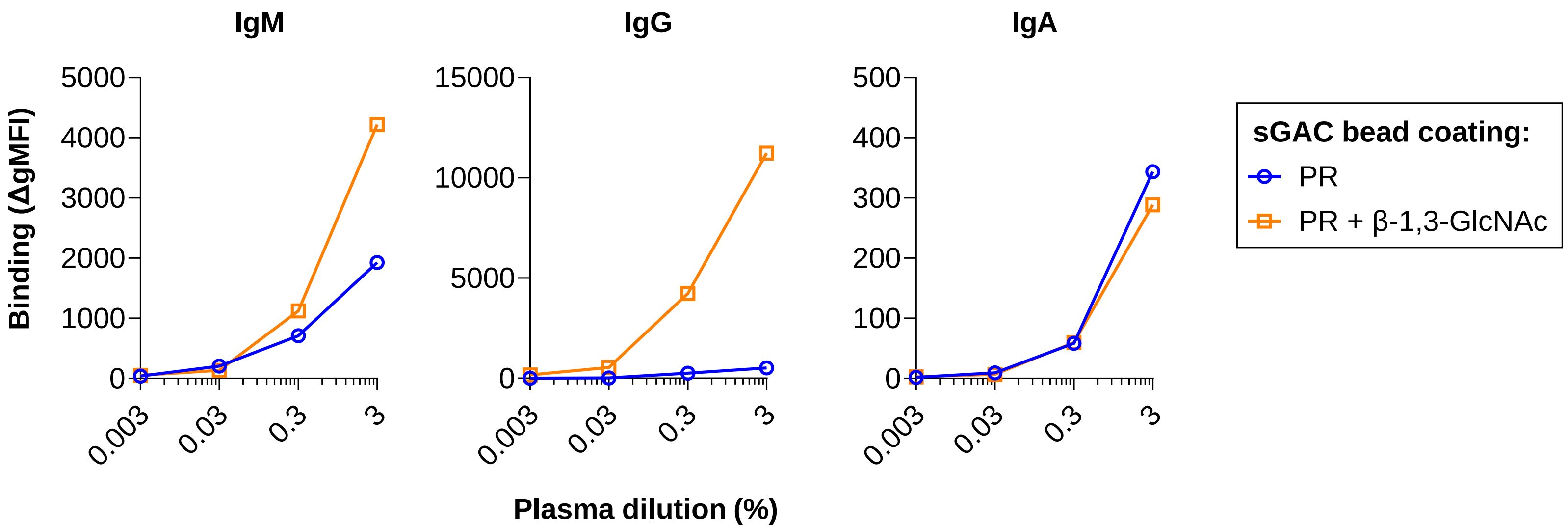

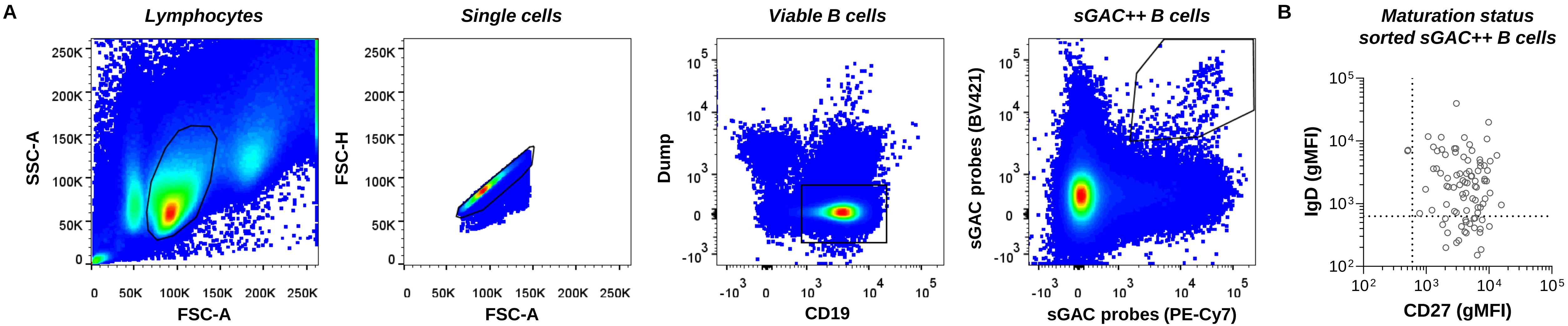

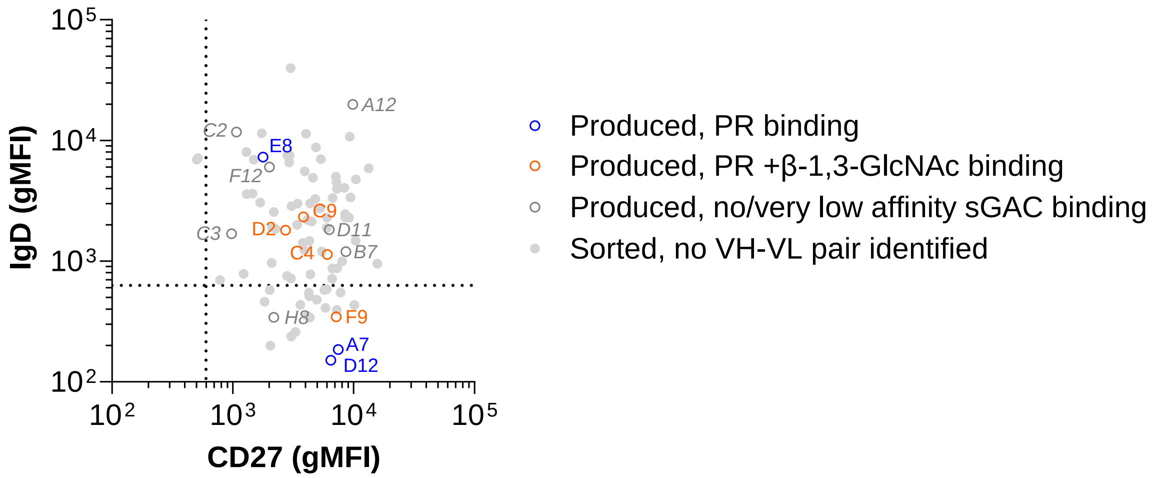

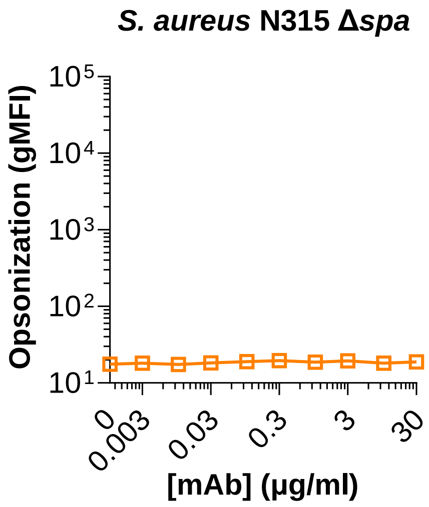
